# Predicting the Yields of Species Occupying a Single Trophic Level with Incomplete Information: Two Approximations Based on the Lotka-Volterra Generalized Equations

**DOI:** 10.1101/2020.12.31.425009

**Authors:** Hugo Fort

## Abstract

The linear Lotka-Volterra generalized equations (LLVGE) serve for describing the dynamics of communities of species connected by negative as well as positive interspecific interactions.

Here we particularize these LLVGE to the case of a single trophic level community with *S* >2 species, either artificial or natural. In this case, by estimating the LLVGE parameters from the yields in monoculture and biculture experiments, the LLVGE are able to produce quite accurate predictions for species yields.

However, a common situation we face is that we don’t know all the parameters appearing in the LLVGE. Indeed, for large values of *S*, only a fraction of the experiments necessary for estimating the model parameters is commonly carried out.

We then analyze which quantitative predictions are possible with an incomplete knowledge of the parameters. We discuss two approximations that allow using these LLVGE as a quantitative tool. First, when we only know a fraction of the model parameters, the mean field approximation allows making predictions on aggregate or average quantities. Second, for cases in which all the interaction parameters involving a particular species are available, we have the focal species approximation for predicting the yield of this focal species.

## 1. THE LOTKA-VOLTERRA GENERALIZED LINEAR MODEL FOR MANY INTERACTING SPECIES

The *Lotka-Volterra equations* (Lotka 1925; Volterra 1926) constitute the classical mathematical model for describing the species abundances in terms of their interactions. Lotka and Volterra derived two different sets of equations: One set applies to situations involving competition for food or space and the other set to the *predator-prey* situation. In fact, the possible interactions between two species are richer and include the following cases:

- mutual competition −/−, each species has an inhibiting effect on the growth of the other;
- amensalism −/0, the growth of one species is negatively affected while the growth of the other is unaffected;
- predation −/+, the ‘predator’ (‘prey’), has an inhibiting (accelerating) effect on the growth of the prey (predator);
- commensalism 0/+, the growth of one species accelerates while the other is unaffected,
- mutual cooperation or mutualism +/+, each species has an accelerating effect on the growth of the other.

[Notice that according to these definitions, the host-parasite and the plant-herbivore interactions would be classified as ‘predation’.]

A unified framework accommodating all kinds of interactions between pair of species are the so-called Lotka–Volterra *generalized* equations (Hofbauer & Sigmund 1998, Pastor 2008, Fort 2020a) Indeed, in general, interspecific interactions different than mutually negative −/interaction between species, like −/+ or +/+ have been considered in the past to occur only between different trophic levels (Morin 2011). For instance, the typical example of −/+ is predation and of +/+ is mutualism, like the plant-pollinator relationship. However, several examples of these types of interspecific interactions have been also found for species sharing the same trophic level; from viruses (Turner & Chao 1999; Arbiza *et al*. 2010) to natural plant communities (Holmgren *et al*. 1997) or artificial plant polycultures (Halty *et al*. 2017), just to mention few examples.

This theory in its simplest formulation consists in a set of linear equations for the per capita growth rates 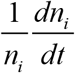 where *n_i_* is a variable denoting the *yield* of species *i* (its density = abundance per unit of area, or its biomass density). A straightforward way to write these *linear* Lotka–Volterra generalized equations (LLVGE) is:

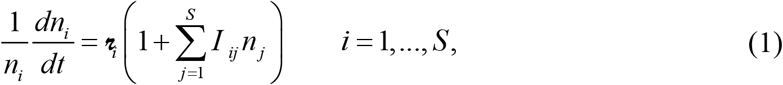

where 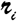 is the intrinsic growth rate of species *i*, with dimension of time^-1^, and *I_ij_* is an interaction coefficient quantifying the *per capita* effect of species *j* on species *i* through their pairwise interaction, which can be either negative (competition), positive (facilitation) or zero, with dimension of density^-1^.

These LLVGE can be thought as the first order or linear approximation in a Taylor series expansion of the per-capita growth rates of species about the equilibrium points of a more complex and general theory (Volterra 1926, 1931):

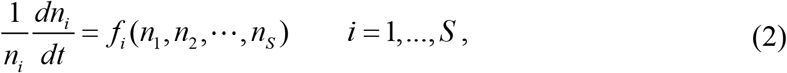

where *f_i_*(*n*_1_, *n*_2_…, *n_S_*) are arbitrary functions of the *S* species making up the community, which we denote more compactly in terms of the *S*-dimensional vector of yields **n** = [*n*_1_, *n*_2_,… *n_S_*] as *f_i_*(**n**). Therefore, let *δ***n** = **n** – **n**\* be an small displacement from an equilibrium point **n**\*, which by definition satisfies *f_i_*(**n**\*) = 0, and if we perform a Taylor expansion of, *f_i_*(**n**) around **n**\* we get:

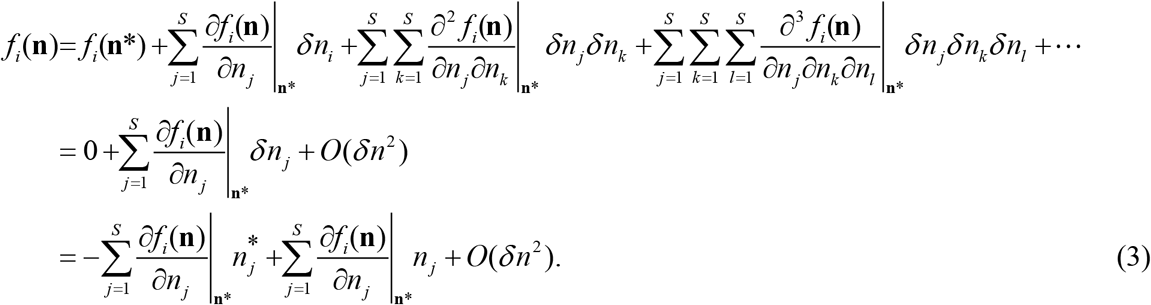

Where *O*(*δn*) involve quadratic terms given by 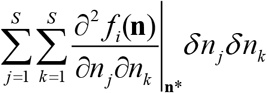, or terms of order greater than two, *i.e*. cubic terms given by 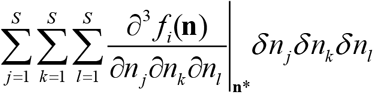, in such a way that we obtain equations (1) by identifying

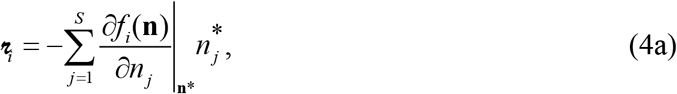

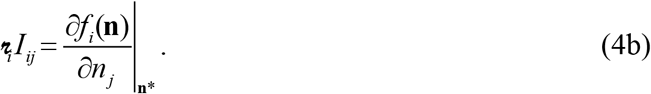

Let us introduce at this point a widely used matrix in community ecology, which is closely related with the matrix [*I_ij_*] but different and that often has been used interchangeably, the *community matrix J_ij_* (Levins 1968). The community matrix, is nothing but the mathematical Jacobian matrix (see Appendix I of Fort 2020a), defined by:

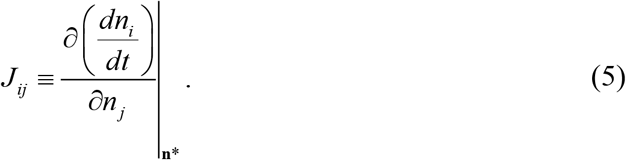

Thus eq. (4b) can be rewritten as connecting the matrix [*I_ij_*] with the community (*i.e*. Jacobian) matrix (see Appendix I of Fort 2020a):

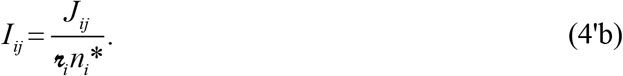

Even though the linear Lotka-Volterra equations have been criticized as being too simple for quantitative modeling real systems involving many interacting species^1^, they can be regarded as ***descriptive*** or ***phenomenological*** models, *i.e*. they describe how the abundance of one species affects the abundance of another, without specifically including a particular mechanism for such interaction (Morin 2011). Neither the nature of the particular competition mechanism nor if it is direct or mediated by other(s) species, is specified. Rather, ***effective*** interaction coefficients summarize the per capita effects of one species on another.

## 2 THE LOTKA-VOLTERRA LINEAR MODEL PARTICULARIZED FOR SINGLE TROPHIC COMMUNITIES

As we mentioned in the previous section, the LLVGE serve to model even single trophic communities. In this section we will consider different versions of single-trophic equations. Firstly, the purely competitive case and secondly the more general case in which species, despite belonging to a single trophic level, can exert besides competition facilitative interactions one on another.

### 2.1 Purely competitive communities

If the effect of all interactions between species has an inhibiting effect on the growth of the other species, i.e. interspecific coefficients are given by:

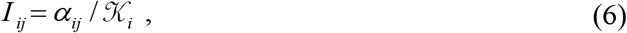

where 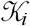 is a maximum sustainable population density, the *carrying capacity*, and the *α_ij_* are negative competition coefficients measuring the per capita effect of species *j* on the abundance of species *i*. It is customary to take the intraspecific competition coefficients *α_ii_* = −1. Hence, we can write the Lotka-Volterra competition equations (LVCE) as:

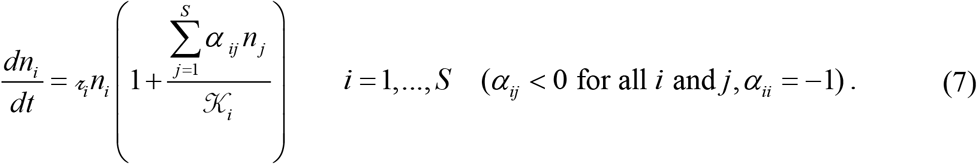

### 2.2 Single trophic communities with interspecific interactions of different signs

The LLVGE for *S* interacting species within a single trophic level can be written as:

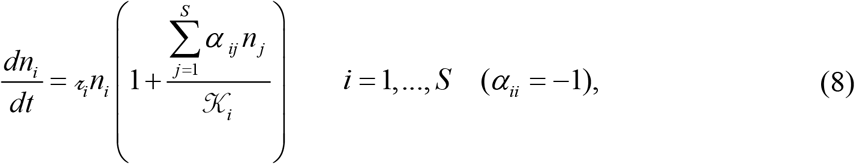

i.e. identical to the competition equations (7) but without the restrictive condition that the interspecific *α_ij_* coefficients be negative.

At this point it is worth summarizing the definitions and interpretations of the three alternative interaction strength matrices we have introduced (Table 1, adapted from Novak et al. 2016).

**Table 1.**
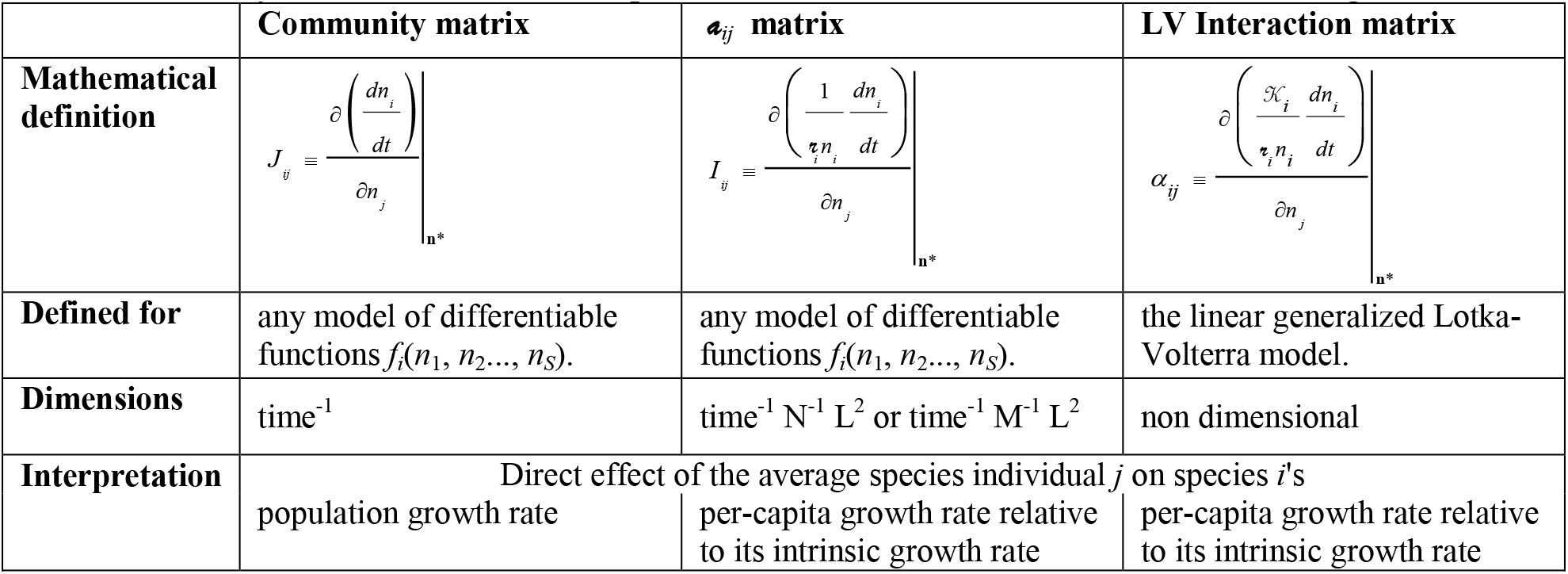
Summary of the definitions & interpretations of the three alternative interaction strength matrices

### 2.3 Obtaining the parameters of the linear Lotka-Volterra generalized model from monoculture and biculture experiments

To obtain the model parameters our starting point are the equilibrium abundances predicted by equation (7) or (8) which verify:

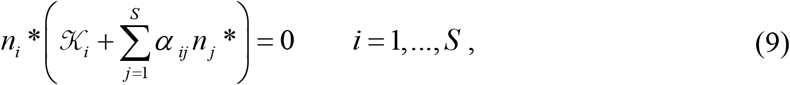

where the asterisks denote quantities at equilibrium. A main difficulty for obtaining from the set of equations (8) the equilibrium species yields 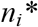 is how to get the set of parameters 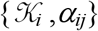. A straightforward procedure is to perform, during sufficiently long enough periods (in order that the equilibrium state is reached): a) the *S* single species or monoculture experiments, and from each of them to estimate the carrying capacities as the yield of the species *i* in monoculture 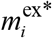 (we use *m* to emphasize that they are yields in monoculture, while the superscript ‘ex’ is for denoting experimentally measured quantities to distinguish them from the theoretical ones); b) the *S*×(*S*-1)/2 pairwise experiments and for each of them, obtain the pair of the *biculture* (pairwise experiments) yields, 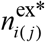 and 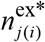(the subscripts *i*(*j*) and *j*(*i*) stand for the relative yield of species *i* in presence of species *j* and *vice ver*sa). Using a) we obtain *K*_i_, and then from b) we obtain 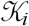, and then from b) obtain *α_ij_* and *α_ji_* by solving equation (2.9) for *S* = 2, as:

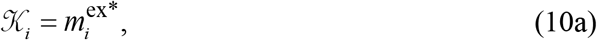

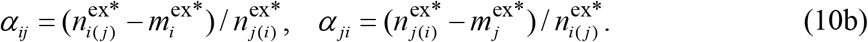

Therefore, if the yield of species *i* in biculture with species *j* is 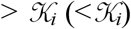 then *α_ij_* > 0 (< 0) and the interaction of *j* on *i* is facilitative (competitive). This is the kind of approach followed by Vandermeer (1969) in a pioneering experimental study with protozoa.

It is worth stressing that the above procedure to obtain the model parameters works by assuming that the community is in a state of equilibrium. In the strict mathematical sense a community is at equilibrium only when the rate of change for all species, i.e. the left hand side of equations (8) is zero. But this theoretical ideal is rarely achieved in natural communities (Wiens 1984; Roxburgh & Wilson 2000). Moreover, in nature the intensity of interactions may vary with a range of environmental factors including climate, kind of resources, spatial distribution of resources and temporal variation in all of the foregoing. Thus, the coefficients themselves become variables.

### 2.4 Quantifying the Accuracy of the Linear Model for Predicting Species Yields

As we have mentioned, the minimality of the LLVGE often raises doubts on their ability to make quantitative predictions, like species abundances. Rather Lotka-Volterra models are regarded more as a qualitative than a quantitative tool in population or community ecology (Brown *et al*. 2001). More ‘realistic’ and complex theories, *e.g*. equations involving non-linear response functions (Vandermeer & Goldberg 2013) and higher order interactions (Abrams 1983) implying non-additive effects (Morin 2011), are often preferred because they are perceived as more reliable (although they can be as intractable as the real systems they aim to model). However, this is done at the price of including additional parameters which are very hard to measure. Actually, in many natural communities –like tropical forests, plankton or mutualistic networks– the species richness *S* is of the order of hundreds. Hence, estimating all the parameters of the LLVGE from empirical data is an unfeasible task, let alone estimating additional parameters of more complex models.

The viewpoint I adopt here is that models should be mainly evaluated not on the basis of the realism of their assumptions but on the basis of the accuracy of their predictions. Thus, in a recent study (Fort 2018b) I analyzed how well the LLVGE work as a *quantitative* tool for explaining/predicting the outcome of 33 experiments for single trophic species belonging to the same taxonomic group (plants, algae, etc.) assuming that a state of equilibrium was reached. With this aim, I used a dataset of experiments designed to measure the effects of intra and interspecific interactions in single-trophic communities with *S* > 2 species, including experimental treatments that measure all the equilibrium yields of:

i. the *S* coexisting species (one treatment),
ii. species in monoculture (*S* treatments),
iii. species in bicultures (*S*×(*S*-1)/2 treatments).

This allowed to obtain the polyculture experimental yields at equilibrium, 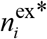, and the monoculture yields 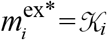, respectively, by taking averages over replicas of (i) and (ii). Similarly, the biculture yields 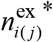 and 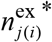 were obtained from replicas of (iii), and the interaction coefficient between species *i* and *j* were in turn computed from 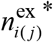 and 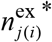 though eq. (10b)^2^.

A commonly used metric to assess how well a model fits observed data is the familiar Pearson’s correlation coefficient (*r*) or its square, the coefficient of determination (*R*^2^). Nevertheless, a problem with these two statistics is that they actually describe the degree of *collinearity* between the observed and model-predicted values rather than their numerical agreement (Willmott 1984). In fact, by their very definition, both indices are insensitive to additive and proportional differences between the model predictions and observations (Willmott 1984). Thus, both suffer from limitations that make them poor measures of model performance. For further discussion on why *r* and *R*^2^ are incorrect measures of predictive accuracy we refer the reader to Li 2017 and references therein. Therefore, to quantitative assess the degree to which the LLVGE match the observations, I resorted to four more appropriate indices (Fort 2018b, Fort 2020a). Indeed, some of these metrics are commonly used in atmospheric and hydrologic sciences. They are:

- The relative mean absolute error (*RMAE*) defined by

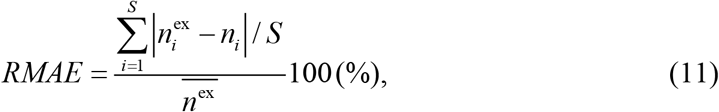

where 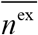 and SE denote, respectively, the mean and standard error for the experimental yields.
- Predictions within 95 % confidence intervals (*P*95), i.e.

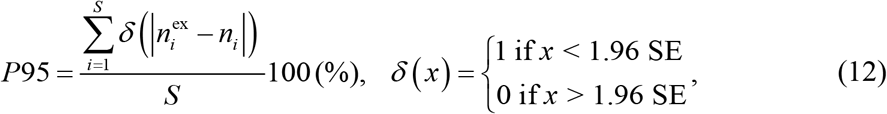
- The modified coefficient of efficiency, defined by

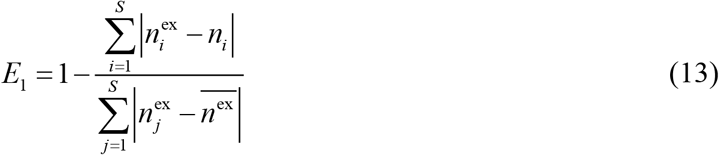
- The modified index of agreement, defined by

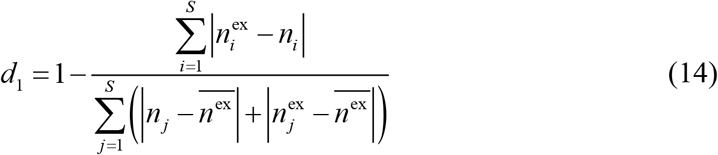

Notice that all the above metrics are in terms of the absolute value instead of the square of differences. This is because absolute values are preferable over squares since by using absolute values errors and differences are given a more appropriate weighting, not inflated by their squared values (Willmott 1981). Squaring in statistics is useful because squares are easier to manipulate mathematically than are absolute values, but use of squares forces an arbitrarily greater influence on the statistic by way of the larger values (Legates & McCabe1999).

Let us briefly comment these indices:

The relative mean absolute error (*RMAE*) is obtained from dividing the mean absolute error (*MAE*) between the mean of the species yields. *MAE* and the similar root mean square error *(RMSE)* are two commonly used measures for assessing the predictive accuracy in the environmental sciences (Li & Heap 2008). To avoid any dependence of *MAE* on *S* we will use the relative metric *RMAE*. In order to quantify the accuracy we need to introduce some reference point. Actually error measures, like *RMAE*, are not accuracy measures, so they can only tell which model produce less error but there are unable to tell how accurate a model is (Li 2017). At any rate, a very tolerant measure of model goodness would be *RMAE* < 100%, *i.e*. every quantity is measured with an error smaller than the size of the quantity itself. We will consider here the more stringent threshold of *RMAE* < 50%. A reference point of 50% might seem too high. Still, it is comparable with the typical SE of the experimental yields (as we shall see in the companion Application 2 chapter).

*P*95 measures the percentage of predictions which fall within the confidence intervals of 1.96*σ* (within the error bars shown in Figure 2). *P*95 = 100 (0) % means that all (none of) the yields predicted by the model fall within the error bars around the corresponding experimental values. For *P*95 we will consider two thresholds: its maximum possible value of 100 % and the (arbitrary) 66.7 %, so that *P*95 ≥ 66.7 % indicates the model does a decent job.

*E*_1_ (Legates & McCabe1999) is a modified version of the coefficient of efficiency (Nash and Sutcliffe 1970) defined by *E*= 1-MSE/*σ*^2^ (*MSE* = mean square error), but in terms of absolute differences rather than square differences. It ranges from minus infinity to 1, the larger its value the better the agreement. In particular, *E*_1_ = 1 indicates perfect match between model predictions and measures. For example, if the absolute differences between the model and the observation is as large as the variability in the observed data (measured by 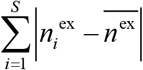), then *E*_1_ = 0.0, and if it exceeds it, then *E*_1_ < 0.0 (*i.e*., the observed mean is a better predictor than *n_i_*). In other words, a value of zero for *E*_1_ indicates that the observed mean 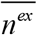 is as good a predictor as the model, while negative values indicate that the observed mean is a better predictor than the model (Wilcox *et al*. 1990).

*d*_1_ (Legates & McCabe1999) is an index similar to *E*_1_. An advantage over *E*_1_ is that it is bounded *i.e*. it varies from 0.0 to 1.0 (again the higher its value the better agreement between model and observations). By the same token it loses the meaningful reference point of 0.0 for the coefficient of efficiency, which serves to assess when the model is a better predictor than the observed mean. However, it is possible to introduce a reference point for *d*_1_ by observing that for two completely uncorrelated random vectors drawn from a uniform distribution, *d*_1_ is on average =1/3 (Fort 2018a, 2020a). Therefore if *d*_1_ ≤ 1/3 we conclude that the model is a poor predictor.

Using the metrics of Table 2 it was shown that the LLVGE can predict with reasonable accuracy not all but the majority of the species yields for most of the experiments.

**Table 2.**
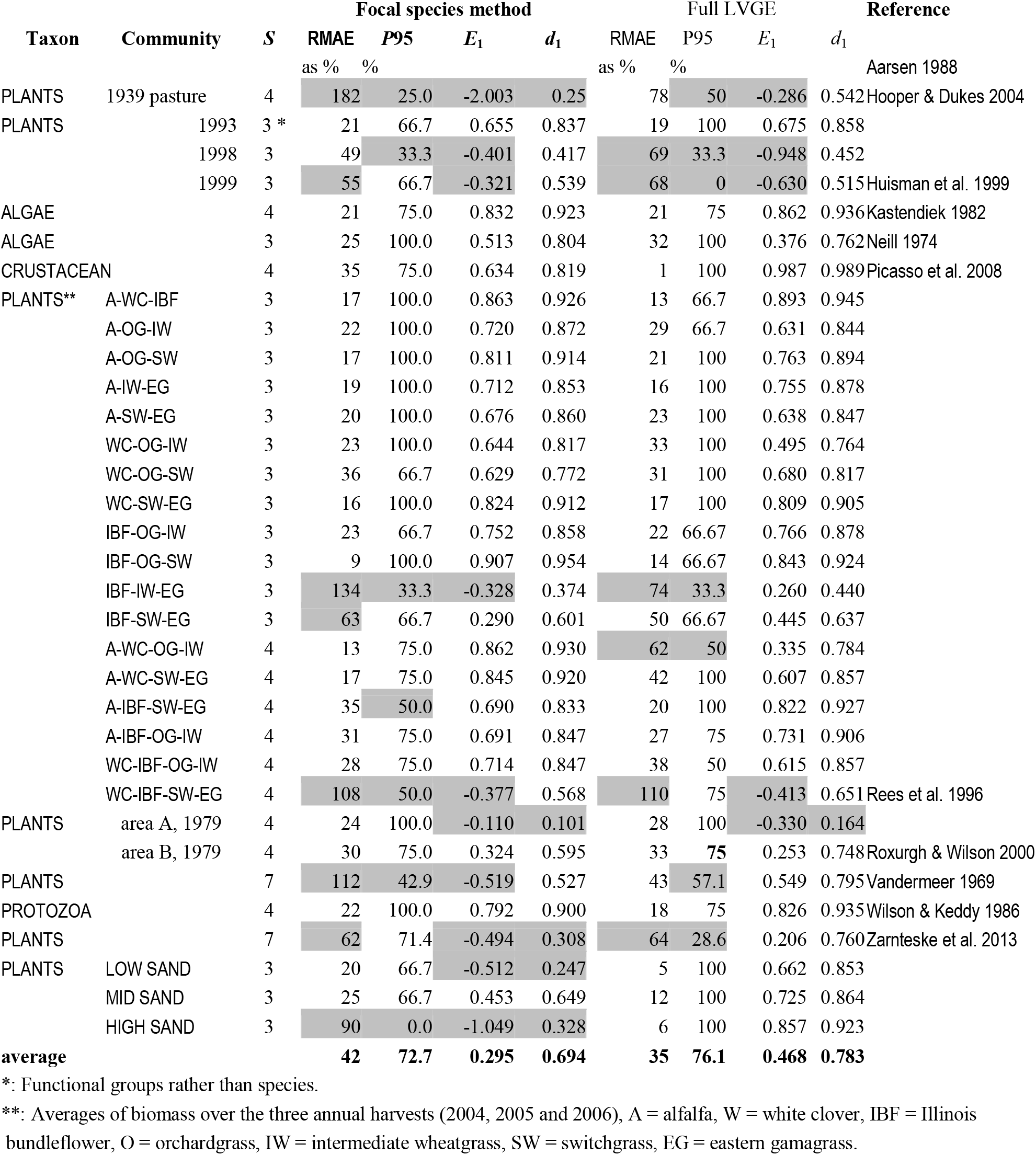
Error and accuracy metrics of the focal method for the 33 experimental studies of Fort (2018a). The first column provides the corresponding references. *S* is the number of species. Columns from *RMAE* % to *d*_1_ are the four metrics to assess model error/accuracy used. Highlighted in gray are values that violate the required accuracy conditions: a) *RMAE* < 50%, b) *P*95 ≥ 66.7%, c) *E*_1_ > 0 and d) *d*_1_ > 1/3. Also listed for comparison the same metrics in the case of using the full interaction matrix as in Fort (2018a).

## 3. WORKING WITH IMPERFECT INFORMATION

A common situation we face when trying to predict yields and final diversity of multi-crops in agriculture or of natural communities is that we don’t know all the parameters appearing in the LLVGE. Indeed, a main limitation of the experimental procedure of performing all the monoculture and biculture treatments is that it is only feasible provided *S* is not too large. This is because the number of required experiments grows as *S*^2^. Thus, for large values of *S*, only a fraction of these experiments is commonly carried out and consequently we have an incomplete knowledge of the *S*^2^ parameters required to compute the equilibrium yields from the LLVGE.

In this section we will consider two situations in which we can still predict some relevant quantities about species yields with an incomplete knowledge of the LLVGE parameters. Both cases correspond to different practical problems in agricultural sciences. To motivate the first case we can think about mixtures of perennial crops grown for biomass, forage, or food production, and the related phenomenon of *overyielding*, i.e. the increased biomass production in species mixtures relative to monoculture (Beckage & Gross 2006). Thus, let us suppose that we only know a fraction of the interaction coefficients. We will show that in this case of incomplete information the LLVGE, via the recently proposed *mean field* approximation (Fort 2018b), can still predict global or aggregate quantities which are important for farmers. The second situation is when one is interested in predicting the abundance of a *particular* species embedded in a community with other species belonging to the same trophic level. Imagine for example a crop coexisting with different species of weeds and that one is interested in designing rational weed management practices. For instance, by enhancing the competitive ability of crops and simultaneously avoiding environmental damage with the use of herbicides (Guglielmini *et al*. 2016). And, let us suppose that for this species we have quite detailed data, like all the interaction coefficients that involve this species of interest. Is it possible to predict the performance of this given particular species≥ We will see that the answer is yes; an approximation that works in such cases in which we know all the parameters corresponding to a single species of particular interest is the *focal species approximation* (Fort 2020a, 2020b).

By performing a bibliographic search, one can check that the number of published experimental studies that measured all the yields (i)-(iii) mentioned at the beginning of section 2.4 decreases quickly as *S* grows. For *S* > 6 is very hard to find an experiment in which the totality of the *S*(*S*+1)/2 treatments were carried out but only a percentage *f_e_* of the them. Nevertheless, in such cases with *f_e_* < 100 %, it is still possible to make quantitative predictions provided we work with relative yields 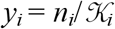(*i.e*. the species yield in mixture normalized by its yield in monoculture) rather than yields *n_i_* and use an approximate expression for the interaction matrix.

It is not difficult to show that the interaction matrix can be written in terms of these relative yields, *y_i_*,

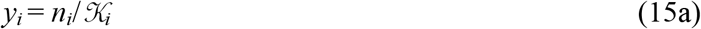

as:

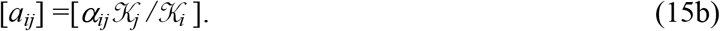

Notice thus that the diagonal elements of the matrix [*a_ij_*], corresponding to intra-specific competition, remain equal to −1 since *a_ii_*=*α_ii_K_i_*/*K_i_* = *α_ii_* = −1. Thus, Eq. (9) for the equilibrium in which all species coexist can be written as (Fort 2020a):

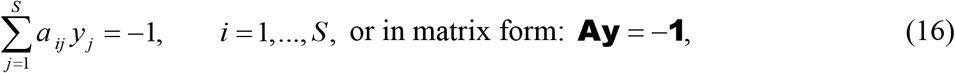

where we use **A** to denote the matrix [*a_ij_*], while **y** and **1** are column vectors of *S* entries with, respectively, relative yields *y_i_* and ones.

So now we are in position to consider the two above mentioned approximations for **A** = [*a_ij_*] which allow making predictions about the yields in cases with incomplete knowledge of the LLVGE parameters.

### 3.1 The *‘Mean Field Matrix’* (MFM) approximation for predicting global or aggregate quantities

The *Mean Field Matrix* (MFM) approximation (Fort 2018b, Fort & Segura 2018), is capable of predicting aggregate or mean quantities, expressed in terms of relative yields, with reasonable accuracy. The name alludes to the resemblance of this approximation with the **Mean Field approximation**, commonly used in physics, consisting in replacing spatially dependent variables by a constant equal to their mean value. Here the average will be taken over the values of the interspecific interaction coefficients *i.e*. the off-diagonal part of the interaction matrix [*a_ij_*] is replaced by a “mean field” competition coefficient.

The aggregate or mean quantities we will consider are the Relative Yield Total (de Wit 1970),

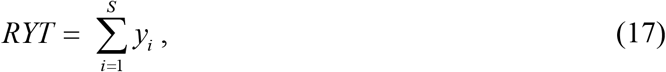

and the Mean Relative Yield,

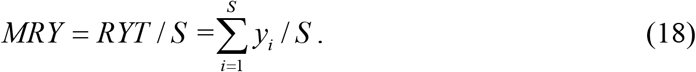

Both these indices allow comparing community productivity on a relative basis. For instance, in agriculture science, the *RYT* is often used to quantify the overyielding of diverse plant mixtures relative to plant monocultures in studies of biodiversity effects on ecosystem function. A *RYT* > 1 implies that the yield performance will be better in polyculture than in monoculture, a phenomenon termed as *overyielding* (Vandermeer 1989).

The idea then is to replace the off-diagonal elements of the interaction matrix [*a_ij_*], corresponding to interspecific interactions, by their average value over the sample of available interspecific competition coefficients:

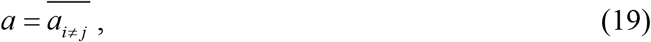

which thereafter will be called the *mean interspecific interaction strength parameter*. In such a way we get a mean field matrix (MFM):

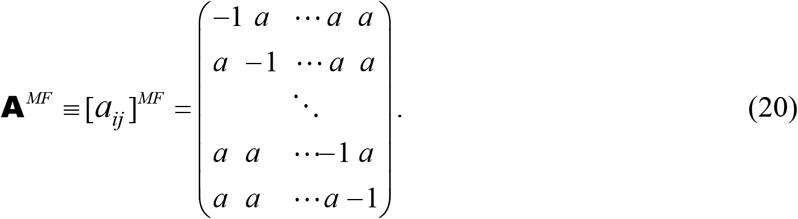

For the MFM Eq. (16) reduces to

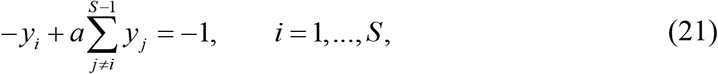

that by summing and subtracting *ay_i_* can be written as:

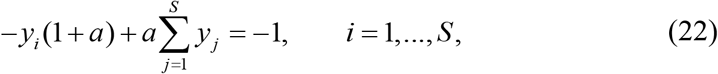

that is,

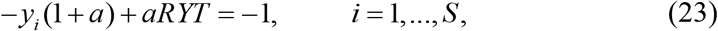

which we can solve for *y_i_* as:

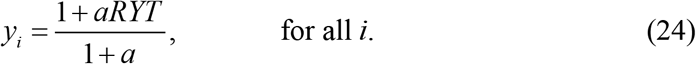

And, if we sum from 1 to *S* both sides of (24), we get

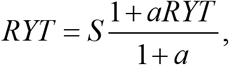

which allows to express the *RYT* and *MRY* as a simple functions of *a* and *S* (Fort 2018b, Fort & Segura 2018):

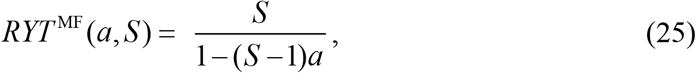

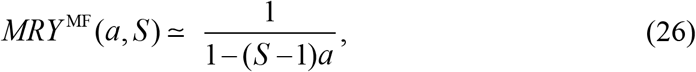

where the superscript ‘MF’ is to emphasize that these expressions hold for the MFM approximation. The pair of Eqs. (25) and (26) tell us that the MFM approximation is particularly suited to address how the total or mean yield depends with the species richness and the mean intensity of competition. This is an interesting question in ecology.

A caveat of the method is that if facilitation is the dominant interspecific interaction, so that *a* > 0, the denominator of equations (25) and (26) could become negative for *a* or *S* sufficiently large. Since the *RYT* and *MRY* must be positive quantities, a requirement for this approximation to work properly is that facilitation is not too large.

Comparisons between predictions generated by formulas (25) and (26) against data from a dataset of 77 experiments (Fort 2018b) are shown, respectively, in Fig. 1 and Fig. 2 as a function of both *S* and *a*. In all these experiments the mean interspecific interaction strength parameter is negative and the number of species *S* varies from 2 to 32 (Fort 2018b).

**Figure 1.**
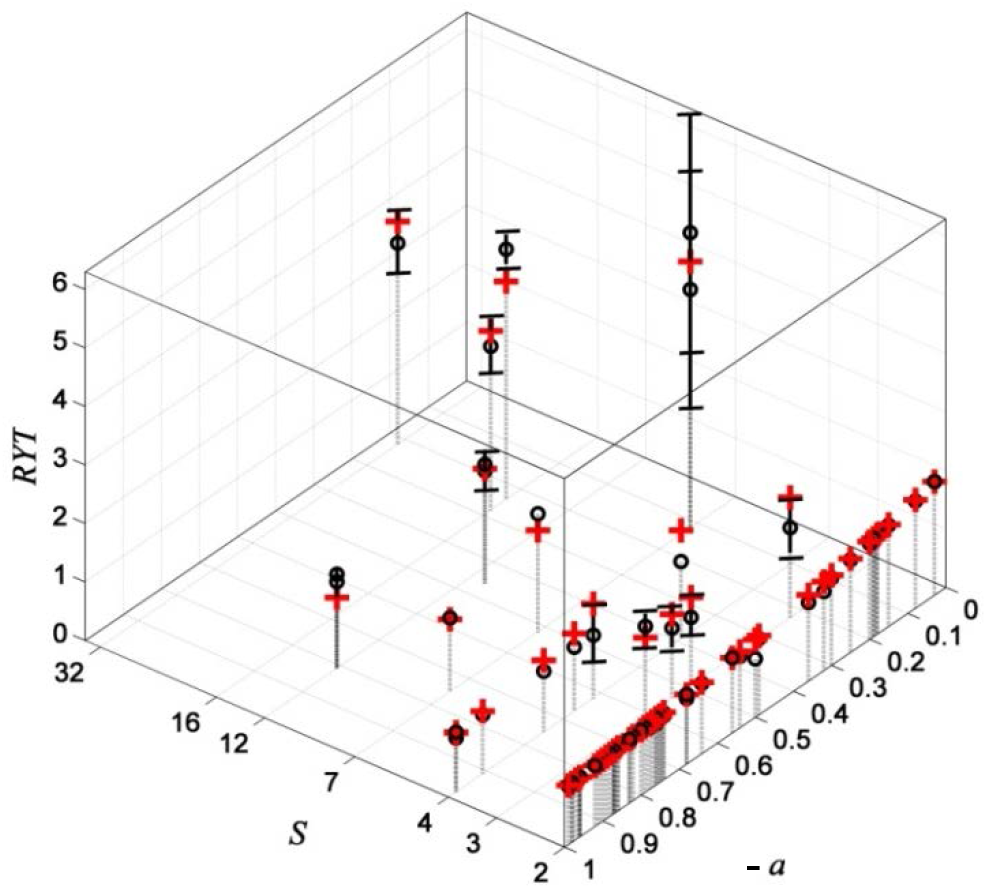
Empirical (o) and theoretical *RYT*, predicted by the MFM approximation Eq. (25) (red crosses), for a set of 77 experiments (see text) as a function of *S* (log scale) and the mean competition parameter *a*. The experimental error bars correspond to ± the standard error (SE).

**Figure 2.**
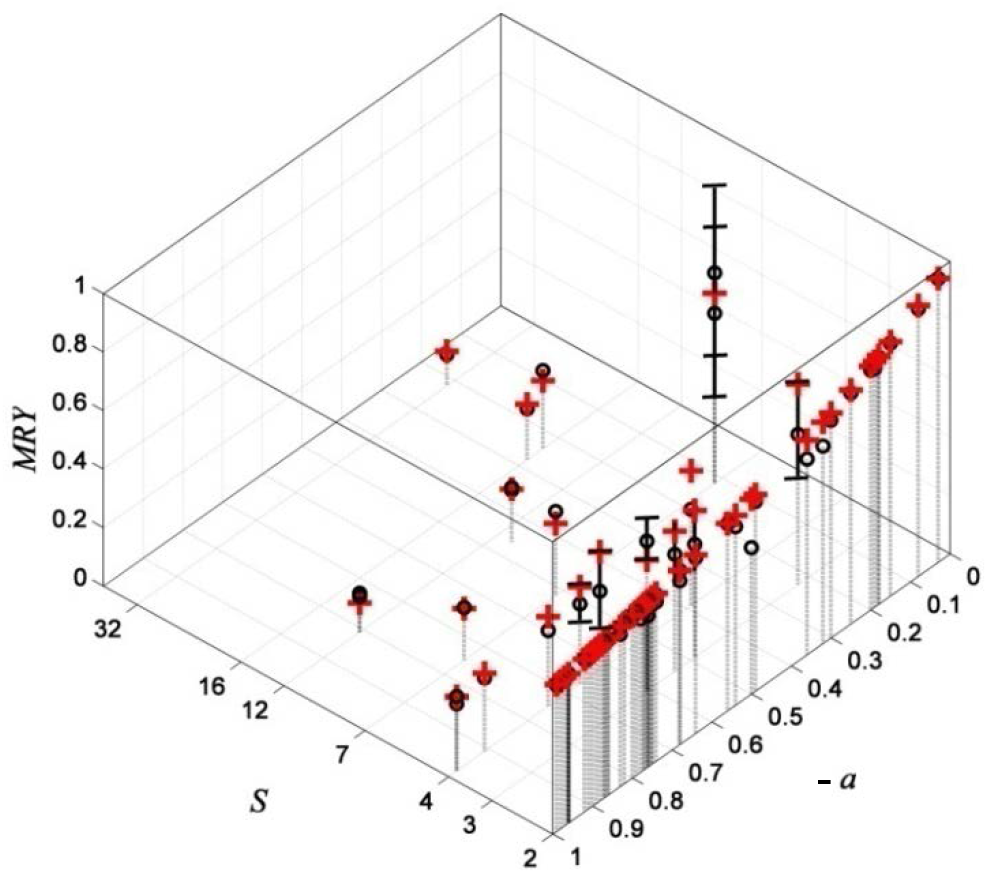
Theoretical *MRY* predicted by the MFM approximation Eq. (26) (red crosses) vs. empirical *MRY* (o) with error bars corresponding to ± SE for a set of 77 experiments as a function of *S*(log scale) and *a*.

As mentioned, the MFM approximation is useful to analyze how the yield depends with the species richness and the mean intensity of competition. Notice that:

- First, regarding the dependence of the *RYT* as a function of *S*, Fig. 1 shows that the *RYT* increases with *S*, a phenomenon which has been observed in many empirical studies and interpreted as *niche partitioning* (Cardinale *et al*. 2011; 2012). That is, different species can use resources in a complementary way and therefore more diverse communities would lead to a higher *RYT*. The *RYT* > 1 for the great majority of the experiments considered here, especially for those with *S* > 2 interacting species (24 out of 25 as shown in Fig. 1), is a clear *quantitative* evidence of such species complementarity (Loreau & Hector 2001, de Wit 1970). Notice that for a fixed value of the mean interaction parameter *a*, Eq. (25) becomes a non-linear saturating relationship between diversity and yield. Non-linear saturating relationships have also been obtained in models by simulation or in other more complex models (Loreau 1998, Schnitzer *et al*. 2011, Downing, et al. 2012).
- Second, as expected, for fixed *S* the *RYT* and *MRY* (Fig. 2) both decrease with the absolute value of *a*. That is, the more intense the average interspecific competition the smaller the yield.
- Third, the relative absolute errors of the *RYT* and *MRY* obtained when using formulas provided by Eq. (25) and Eq. (26), in the case of experiments with incomplete knowledge of model parameters, range between 0.8% and 24 % (mean 12.4 % ± 4.6 %) and are not significantly greater than those obtained by just solving the entire set of LLVGE, which was possible for experiments in which all the model parameters were available as the ones considered in Fort (2018a). Furthermore, in general the *RMAE* is proportional to the size of the experimental error bar and thus the failure of equations (25) and (26) can be mostly explained by a large variance among experimental replicas or by a low experimental precision.

Therefore:

**Provided only competition interactions among species exist or at least they are dominant, we still can predict *aggregate* (*RYT*) or *mean* (*MRY*) relative competition indices with an accuracy inversely proportional to the experimental variance of the experimental measurements.**

### 3.2 The *‘Focal species’* approximation for predicting the performance of a given species when our knowledge on the set of parameters is incomplete

Now, as mentioned at the beginning of this section, let us turn to a different situation and suppose we are interested in the performance of a particular or *focal* species that coexists with other species. For example, say a crop that coexists with several species of weeds. Since we are not interested in the yield of each individual weed, *we can treat all of them as a single species*. Thus in order to make more accurate specific predictions for this focal species than just the *MRY*, one possibility is to measure the interaction coefficients for this species *k* with all the others and use a *focal interaction matrix*, in terms of relative yields, of the form:

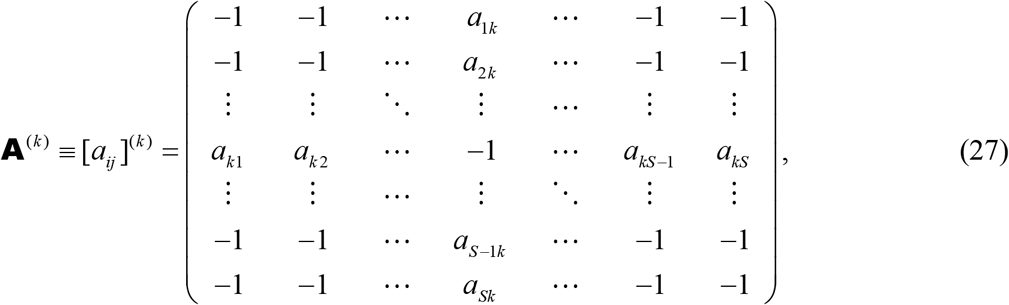

*i.e*. all the interspecific coefficients between species different from the focal species *k* are taken equal to −1, while *a_ik_* denotes the interaction coefficient of the focal species over species *i* and *a_ik_* the interaction coefficient of species *i* over the focal one.

The recipe is then to use matrix (27) to solve Eq. (16) and only take into account the relative yield for the focal species 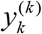 (while neglecting the values 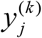 for *j* ≠ *k*). That is, from a matrix of the form (27) we obtain just one relative yield for the focal species 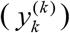. Let us call this approximation, which represents and improvement of the mean field one, the *focal species* approximation (Fort 2020b).

Obtaining the 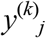 would be straightforward from equations if A) the theoretical coexisting equilibrium were *feasible, i.e*. 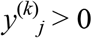 *for all species* (to warrant that Eq. (9) reduces to Eq. (16)), and B) the matrix **A**^(*k*)^ were invertible, by simple inversion of the matrix equation between parentheses in equation (16). However, it is easy for *S* > 3 species to prove that the focal matrix is non-invertible: the det **A**^(*k*)^ = 0. This determinant can be immediately computed by expanding the determinant of matrix (4) along its focal row or column. For *S* = 3 species, there is generally only one row or column with all elements equal to −1 and the determinant is not necessarily zero. However, in this case it is easy to show that at least one of the species must become extinct at equilibrium (Fort 2020b), and thus condition A) is not fulfilled, meaning that the equilibrium state with all species coexisting is unfeasible *i.e*. 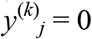 for some species *j*.

Therefore, since either conditions A) or B) are violated, one cannot obtain the equilibrium relative yields by inverting Eq. (16) and has to obtain the equilibria of focal matrices by numerically integrating the dynamical equations:

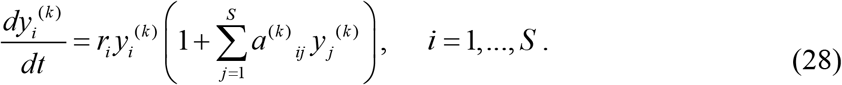

To do this, one can use either a simple finite differences method or an ordinary differential equation solver. Once the set of *S* relative yields 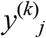 is obtained, the next step is to consider the relative yield for the focal species 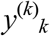 (neglecting the values 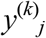 for *j* ≠ *k*).

To illustrate how this approximation works we will consider the experiment of four algal species of Huisman *et al*. (1999): *Aphanizomenon flos-aquae* (*Ap*, species #1), *Chlorella vulgaris* (*Ch*, species #2) a *Microcystis* strain (*Mi*, species #3) and and *Scenedesmus protuberans* (*Se*, species #3) and. The population densities, measured as millions per mL) of the four species in polyculture were:

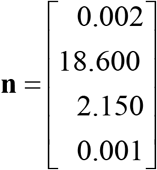

The carrying capacity vector **k** (millions per mL) and the interaction matrix **α** = [*α_ij_*] are given by (Fort 2018a):

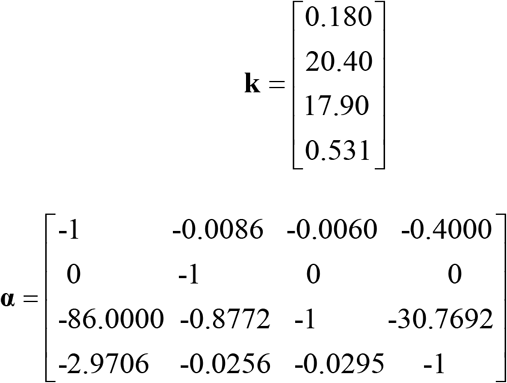

Therefore, using respectively Eq. (15a) and (15b), we get for the vector of experimental relative yields **y**^ex^ and for the matrix **A** = [*a_ij_*]:

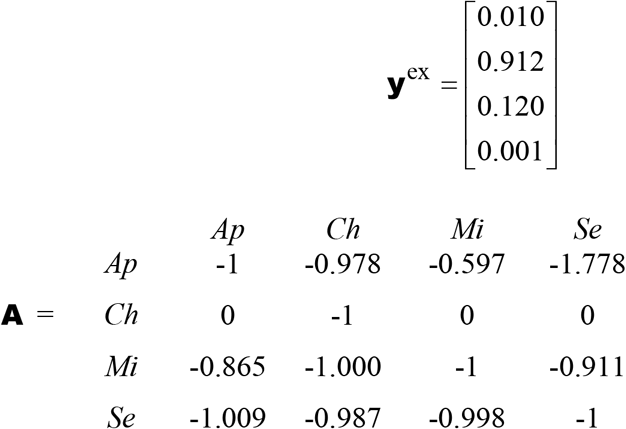

If we consider that the focal species is for example *Ch*, then we obtain the focal matrix 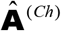 (with boxed elements unchanged):

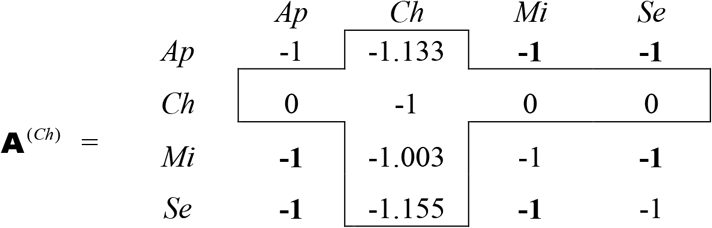

The matrix **A**^(*Ch*)^ is not invertible (three columns are equal and thus its determinant is 0). Therefore we cannot use Eq. (16) to get the equilibrium vector **y**. Nevertheless we can solve the differential equations (28) numerically, until the equilibrium state is reached, and we obtain:

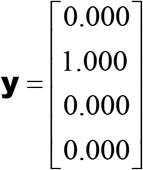

Thus 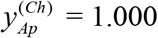, differs in 10 % respect to the experimental 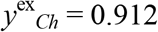 (second entry of Eq. (31)). This is not so bad. Similarly, one can check that 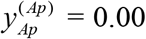 slightly differs with respect to its experimental value of 0.01.

Table 2 summarizes the *RMAE* and the three accuracy metrics −*P*95, *E*_1_ and *d*_1_. It shows highlighted in gray when the indices are below thresholds, indicating poor accuracy.

The RMAE when using the focal interaction matrix and the full interaction matrix are very similar, though P95 and E1 indices show a greater number of experiments below the established accuracy value for the species focal approach. Overall, the accuracy shown for the proposed focal species approximation is similar to that of LLVGE using the full interaction matrix.

Figure 3 compares the results of using this approximation (open big circles) with those when using the full set of LLVGE (filled small circles) for the same four experiments shown in Fig. 2.3. We can see that the accuracy with which the focal species approximation predicts the focal species is comparable to the one obtained when using the full set of LLVGE.

**Figure 3.**
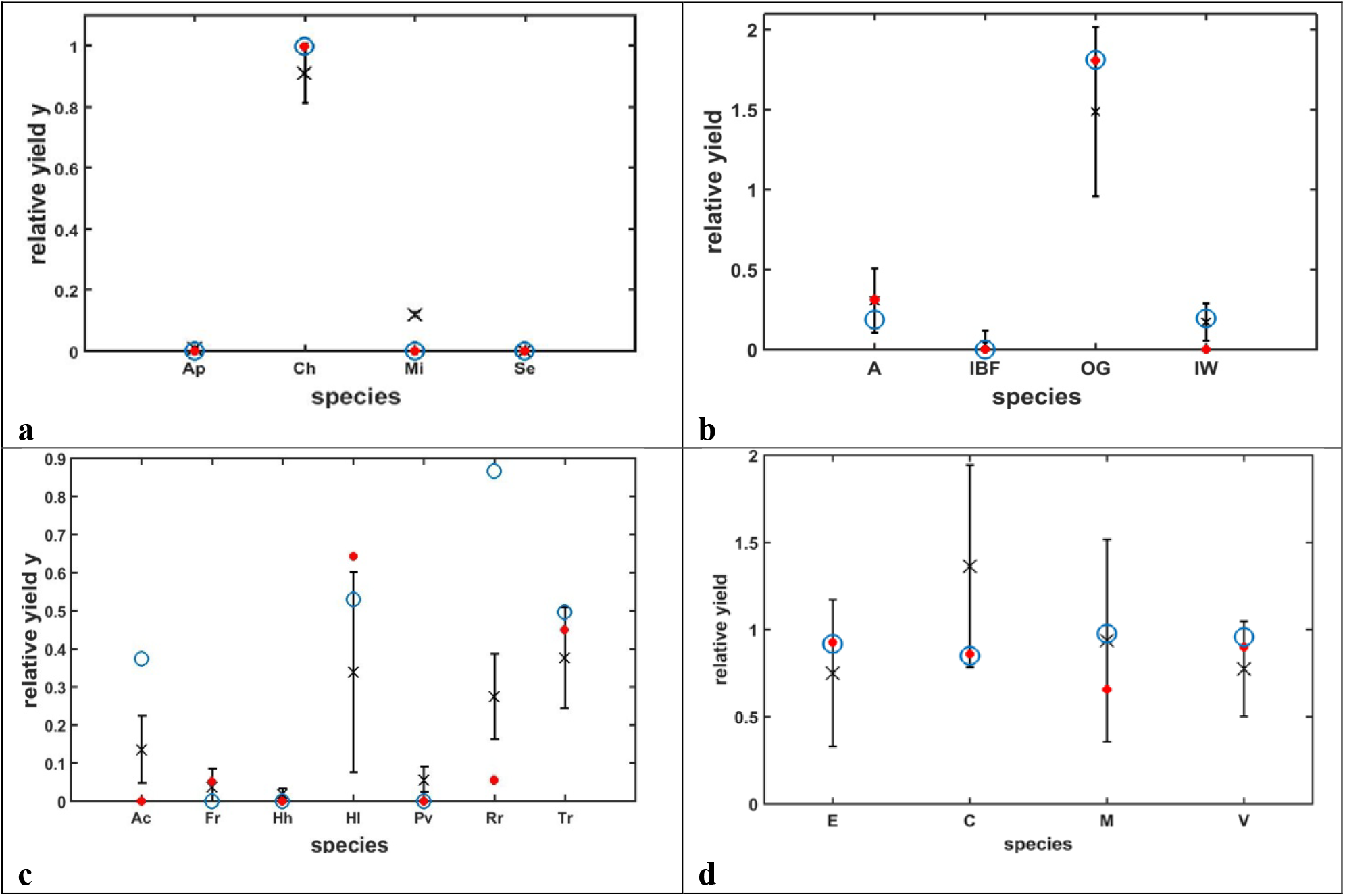
The focal species approximation compared with the full set of LLVGE. Experimental relative yields *R_i_* = *Y_i_/K_i_* with their error bars (crosses), predicted values ex ante by the LLVGE (red filled circles) and the corresponding values produced by the focal species approximation (open blue circles): a) Algae (Huisman *et al*. 1999). b) Plants A-IBF-OG-IW mixture (Picasso et al. 2008). c) Plants (Roxburgh & Wilson 2000). d) Plants, Area A (Rees *et al*., 1996).

Therefore we conclude that:

**It is possible to predict the yield of a given particular species when we just know the interaction coefficients involving this focal species, by means of the focal species approximation, with accuracy comparable to the one obtained when using the full set of LLVGE.**

## 4 CONCLUSION

The linear Lotka-Volterra generalized equation (LLVGE) are the simplest system of equations to model a community of species with all possible types of interspecific interactions: −/−, −/0, −/+, +/0 and +/+. Particular cases of such equations are the competition model (all species interactions −/−) and the mutualistic model (all species interactions +/+).

These LLVGE can be used as a quantitative tool. If all the model parameters are known, the full set of LLVGE allows in general to predict the yields of all species making up the community with reasonable accuracy (Fort 2018a, 2020a). More elaborate models may be necessary for describing and predicting species abundances for particularly complex communities (Brown *et al*. 2001). However, it seems that such cases would be more the exception than the rule (Fort 2018a, 2020a).

However, we have also seen that for the majority of real communities we only have access to a fraction of the model parameters. In these circumstances with incomplete information, we still can make quantitative predictions. Firstly, we have the mean field approximation (MFA). The MFA allows making predictions on aggregate or average quantities, involving the whole community of coexisting species. This has a direct application in agriculture to predict metrics that serve to quantify the performance of mixtures of crops grown for biomass, forage, or food production. Secondly, for such cases in which all the interaction parameters involving a particular species are available, we have the focal species approximation which can be used for predicting the yield of this focal species. Again, agriculture offers an example of a field in which this approximation can be used in the form of a crop coexisting with several species of weeds.

## Footnotes

1 For example, for *S* > 2 species the model would be valid only to the extent that higher-order interactions are null or negligible compared to pairwise interactions (Simberloff 1982). When a pair of species is entangled within a large and complex web of interacting species, like in food webs, it is possible to generate interactions with third or fourth parties that create mutually negative effects for the pair under consideration. This phenomenon is called ‘apparent competition’ (Keddy 2001).

2 Thereafter, since we always assume an equilibrium or quasi-equilibrium state, we will omit the * for all yields in the understanding that they are yields in equilibrium.

## REFERENCES

Aarssen, L.W. 1988. Pecking order of species from pastures of different ages. Oikos 51, 3–12.

Abrams, P. A. 1983. Arguments in favor of higher order interactions. Am. Nat. 121, 887–891.

Arbiza, J., Mirazo, S. and Fort, H. 2010. Viral quasispecies profiles as the result of the interplay of competition and cooperation, BMC Evolutionary Biology 2010, 10:137.

Beckage, B., and Gross, L. J. 2006. Overyielding and species diversity: what should we expect? New Phytologist, 172, 140–148.

Brown, J.H., T.G. Whitham, S.K.M. Ernest, C.A. Gehring. 2001 Complex species interactions and the dynamics of ecological systems: long-term experiments. Science 293, 643–650.

Burton, T. A. 1969 On the construction of Lyapunov functions. SIAM J. Appl. Math. 17 1078–1085.

Cardinale, B. J. et al. 2012. Biodiversity loss and its impact on humanity. Nature 486, 59–67.

Cardinale, B. J. et al. 2011. The functional role of producer diversity in ecosystems. Am. Jour. of Bot. 98, 572–592.

Downing, et al. 2012. The resilience and resistance of an ecosystem to a collapse of diversity. PloS one 7, e46135.

Fort, H. 2020a Ecological Modelling and Ecophysics: Agricultural and environmental applications (IOP ebooks), IOP, Bristol, UK.

Fort, H. 2020b Making quantitative predictions on the yield of a species immersed in a multispecies community: The focal species method. Ecological Modelling 430, 109108.

Fort, H. 2018a On predicting species yields in multispecies communities: Quantifying the accuracy of the linear Lotka-Volterra generalized model. Ecological Modelling 387, 154–162.

Fort, H. 2018b Quantitative predictions from competition theory with an incomplete knowledge of model parameters tested against experiments across diverse taxa. Ecological Modelling 368, 104–110.

Fort, H. & Segura, A. 2018 Competition across diverse taxa: quantitative integration of theory and empirical research using global indices of competition. Oikos 127, 392–402.

Goh, B. S. 1977 Global stability in many species systems. The American Naturalist, 111, 135–143.

Goh, B. S. 1980 Management and analysis of biological populations. Elsevier, Amsterdam.

Guglielmini, A.C., Verdú, A.M.C., Satorre, E.H. 2017 Competitive ability of five common weed species in competition with soybean. Int. J. of Pest Management 63, 30–36.

Gurel, O. and Lapidus, L. 1968 Stability via Liapunov’s second method. Ind. Eng. Chem. 60, 1–26.

Hafstein, S. F. 2007 An algorithm for constructing Lyapunov functions. Electronic Journal of Differential Equations, Monograph 08. URL: http://ejde.math.txstate.edu or http://ejde.math.unt.edu

Halty, V. et al. 2017 Modelling plant interspecific interactions from experiments of perennial crop mixtures to predict optimal combinations, Ecological Applications 27, 2277–2289.

Hector, A. et al. 2010 Ecological Archives E091-155-S1. http://www.esapubs.org/archive/ecol/E091/155/

Hofbauer M. and Sigmund, K. 1998 Evolutionary Games and Population Dynamics, Cambridge University Press, Cambridge, UK.

Holmgren, M., Scheffer, M. and Huston, M.A. 1997. The interplay of facilitation and competition in plant communities. Ecology 78, 1966–1975.

Hooper, D. U. & Dukes, J. S. 2004 Overyielding among plant functional groups in a long-term experiment Ecol. Lett. 7, 95–105.

Hubbel, S.P. 2002 The Unified Neutral Theory of Biodiversity and Biogeography. Princeton Univ. Press, Princeton, N.J.

Huisman, J., Jonker, R.R., Zonneveld, C. & Weissing, F.J. 1999 Competition for light between phytoplankton species: experimental tests of mechanistic theory. Ecology 80, 211–222.

Kastendiek, J. 1982 Competitor-mediated coexistence: interactions among three species of macroalgae. J. Exp. Mar. Biol. Ecol., 62, 201–210.

Keddy, P.A. 2001. Competition (2nd edition). Kluwer Academic Publishers.

LaSalle, J. and Lefschetz, S. 1961 Stability by Liapunov’s Direct Method. Academic Press, New York, N.Y.

Legates, D. R. & McCabe, G. J. 1999 Evaluating the use of “goodness-of-fit” measures in hydrologic and hydroclimatic model validation. Water Resources Research, 35, 233–241.

Levins, R. 1968 Evolution in changing environments: Some theoretical explorations. Princeton Unive. Press, Princeton, NJ. _2_

Li, J. 2017 Assessing the accuracy of predictive models for numerical data: Not *r* nor *r*^2^, why not? Then what? PLoS ONE 12(8): e0183250.

Li, J. & Heap, A. 2008 A Review of Spatial Interpolation Methods for Environmental Scientists. Geoscience Australia, Record 2008/23, 137pp.

Loreau, M. 1998. Biodiversity and ecosystem functioning: A mechanistic model. PNAS 95, 5632–5636.

Loreau, M, & Hector, A. 2001 Partitioning selection and complementarity in biodiversity experiments. Nature 412, 72–76.

Lotka, A. J. 1925 Elements of Physical Biology. Williams and Wilkins, Baltimore.

May, R. M. 1974 Stability and Complexity in Model Ecosystems. Princeton University, Princeton, NJ.

Morin, P.J. 2011 Community Ecology. Wiley-Blackwell, Chichester, UK.

Nash, J. E., and Sutcliffe, J. V. 1970 River flow forecasting through conceptual models, I, A discussion of principles. J. Hydrol., 10, 282–290.

Neill, W.E. 1974 The Community Matrix and Interdependence of the competition coefficients. Am. Nat. 108, 399–408.

Novak, M. et al. 2016 Characterizing species interactions to understand press perturbations: What is the community matrix? Annu. Rev. Ecol. Evol. Syst. 47, 409–432.

Pastor, J. 2008 Mathematical ecology of populations and ecosystems. Wiley-Blackwell.

Picasso, V., Brummer, E. C., Liebman, M., Dixon, P. M. & Wilsey, B. J. 2008 Crop species diversity affects productivity and weed suppression in perennial polycultures under two management strategies. Crop. Sci. 48, 331.

Rees, M.P. et al. 1996. Quantifying the impact of competition and spatial heterogeneity on the structure and dynamics of a four-species guild of winter annuals. Am.Nat. 147, 1–32.

Roxburgh, S.H. & Wilson, J.B. 2000 Stability and coexistence in a lawn community: mathematical prediction of stability using a community matrix with parameters derived from competition experiments. Oikos 88, 395–408.

Schnitzer et al. 2011 Soil microbes drive the classic plant diversity-productivity pattern. Ecology 92, 296–303.

Schultz, D.G. 1965 The generation o f Liapunov functions. In: C T. Leondes (Editor), Advances in Control Systems, Vol. 2. Academic Press, New York, N.Y., pp. 1—64.

Simberloff, D. 1982 The status of competition theory in ecology. Ann. Zool. Fenn., 19, 241–253.

Strogatz, S. H. 1994 Nonlinear dynamics and chaos. With Applications to Physics, Biology, Chemistry, and Engineering Perseus Books, Reading, Massachusetts.

Turner, P. E. and Chao, L. 1999 Prisoner’s dilemma in an RNA virus. Nature 398, 441–443.

Vandermeer, J.H. 1969 The competitive structure of communities: an experimental approach with protozoa. Ecology, 50, 362–371.

Vandermeer, J.H. 1989 The Ecology of Intercropping. Cambridge University Press, Cambridge.

Vandermeer, J. H., & Goldberg, D. E. 2013 Population ecology: First principles. 2nd Edition. Princeton: Princeton University Press.

Volterra, V. 1931 Leçons sur la Théorie Mathématique de la Lutte pour la Vie. Gauthier-Villars, Paris.

Volterra, V. 1926 Fluctuations in the abundance of a species considered mathematically. Nature 118, 558–560.

Wiens, J. A. 1984 On understanding a non-equilibrium world: myth and reality in community patterns and processes. In: Ecological communities: conceptual issues and the evidence. Princeton Univ. Press, Princeton, NJ, pp. 439–457.

Wilcox, B. P., Rawls, W. J., Brakensiek, D. L. and Wight, J. R. 1990 Predicting runoff from rangeland catchments A: comparison of two models, Water Resour. Res, 26, 2401–2410.

Willems, J. L. 1970 Stability Theory of Dynamical Systems. Nelson, London.

Willmott, C. J. 1984 On the evaluation of model performance in physical geography, in Spatial Statistics and Models, edited by G. L. Gaile, and C. J. Willmott, pp. 443–460, D. Reidel, Norwell, Mass.

Willmott, C. J. 1981 On the validation of models, Phys. Geogr., 2, 184–194.

Wilson, S. D. & Keddy, P. A. 1986 Species competitive ability and position along a natural stress/disturbance gradient. Ecology, 67, 1236–1242.

de Wit, C. T. 1970 Proc. Adv. Study Inst Dyn. Numbers Popul. 269.

Zarnetske P. L., Gouhier T. C., Hacker S. D., Seabloom E.W., Bokil V. A. 2013 Indirect effects and facilitation among native and non-native species promote invasion success along an environmental stress gradient. Journal of Ecology, 101, 905–915

